# *RFC1* AAGGG pentanucleotide repeats form parallel G-quadruplex: structural implications for aberrant molecular cascades in CANVAS

**DOI:** 10.1101/2023.08.06.552146

**Authors:** Yang Wang, Junyan Wang, Zhenzhen Yan, Jianing Hou, Liqi Wan, Jie Yi, Pei Guo, Da Han

## Abstract

A pentanucleotide repeat expansion (PRE) of (AAGGG)_n_ in the replication factor C subunit 1 (*RFC1*) gene is recently identified as the genetic cause of cerebellar ataxia, neuropathy, and vestibular areflexia syndrome (CANVAS), and also linked to several other neurodegenerative disorders including the Parkinson’s disease. However, the molecular mechanism by which *RFC1* PRE drives pathology remains poorly understood. Here, for the first time we discovered and determined the high-resolution structures of parallel G-quadruplex formed by AAGGG repeats within the pathogenic *RFC1* PRE, revealing an intriguing conformational plasticity at the 3’-termi that allows stacking of multiple G4s. We further identify a molecular mechanism by which the DNA G4 in *RFC1* PRE impedes polymerase processivity leading to replication stalling and transcription inhibition *in vitro* in a repeat-length-dependent manner, and the transcription inhibition could partially contribute to a reduced gene expression in cells. Our results demonstrate that the DNA G-quadruplex of *RFC1* PRE participate in aberrant molecular cascades, and provide an unprecedented high-resolution structural target to discover helicases and ligands that resolve the pathogenic G4 for therapeutic intervention.

## Introduction

Nucleotide repeat expansion disorders (REDs) constitute some of the most common inherited neurodegenerative diseases, including the Huntington’s disease, amyotrophic lateral sclerosis and frontotemporal dementia (ALS/FTD)^1–3^. Intriguingly, REDs manifest with sequence specificity and only when the number of repeats in the mutational gene exceeds a certain threshold. Recently, a pentanucleotide repeat expansion (PRE) of (AAGGG)_n_ in a noncoding region of the replication factor C subunit 1 (*RFC1*) gene has been linked to cerebellar ataxia, neuropathy, and vestibular areflexia syndrome (CANVAS), which is a neurological disorder of autosomal recessive inheritance characterized by adult-onset and slowly progressive sensory neuropathy, cerebellar dysfunction and vestibular impairment^4–7^.

Normal human *RFC1* intron 2 has a highly polymorphic benign configuration in the polyA tail of an Alu element (AluSx3), including predominantly (AAAAG)_11_, and rarely PRE of (AAAAG)_exp_ and (AAAGG)_exp_ (Fig. 1a). However, *RFC1* PRE of (AAGGG)_exp_, ranging from 400 to 2000 repeats, is found to be a pathogenic and causative factor for CANVAS. *RFC1* serves as a vital gene encoding the large subunit of replication factor C, which is a key and essential DNA polymerase accessory protein required for DNA replication and repair in human cells^8^. Meanwhile, the AluSx3 is known to function in gene regulations via binding proteins to its polyA sequence^9^. It is anticipated that the pathogenic (AAGGG)_exp_ perturbs the AluSx3 activity and causes RFC1 protein loss-of function^4–7^. Two latest works have reported a reduction of full-length RFC1 protein level in CANVAS patients and thus suggested a potential mechanism of protein loss-of-function^10, 11^.

**Fig. 1.**
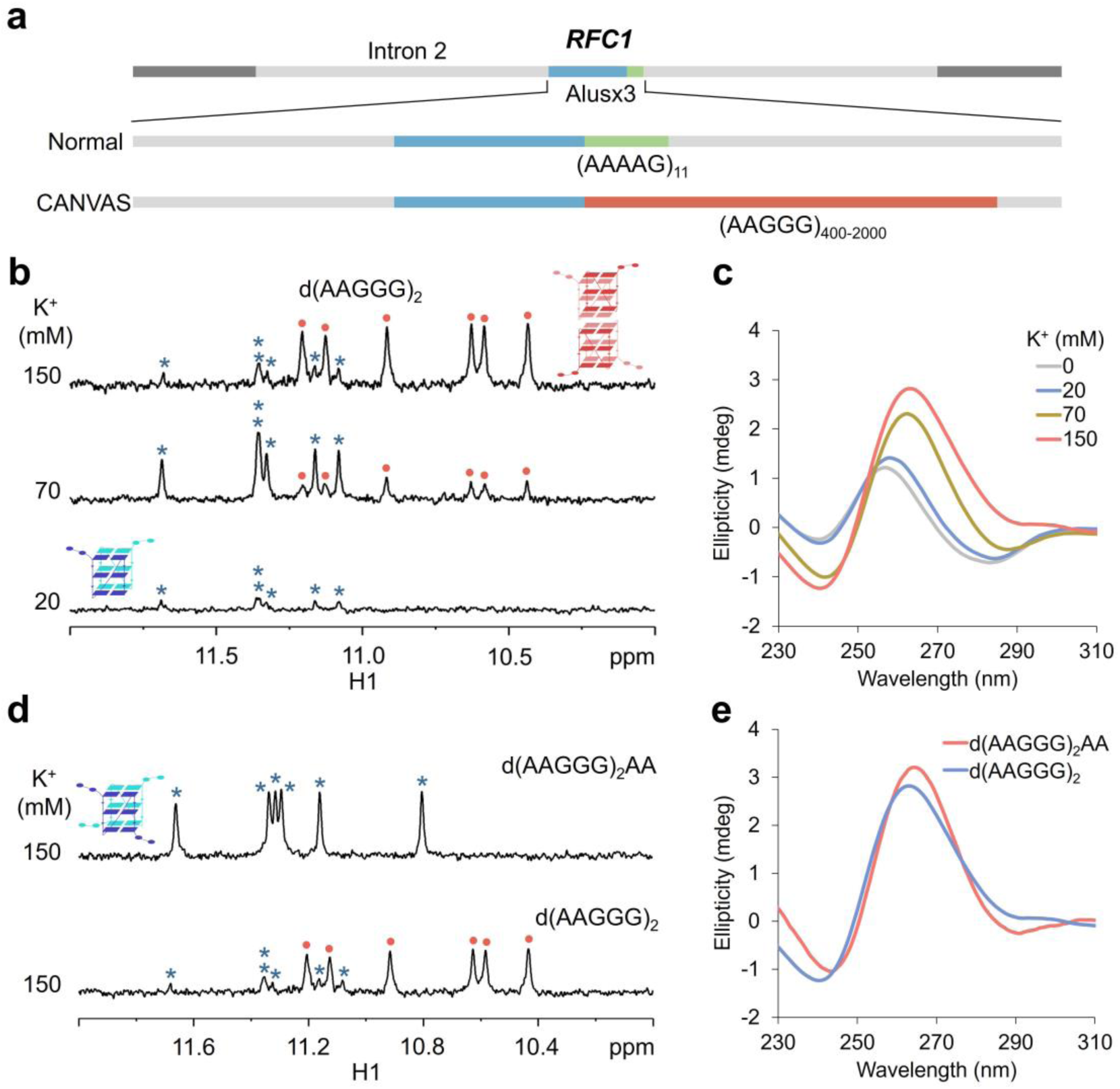
Formation of DNA G4 in the pathogenic *RFC1* AAGGG repeats. **a** Schematic of the reference *RFC1* allele in normal individuals and pathogenic *RFC1* AAGGG repeat expansion in CANVAS patients. **b-e** 1D ^1^H NMR and CD spectra of d(AAGGG)_2_, d(AAGGG)_2_A and d(AAGGG)_2_AA in K^+^ solutions.

As one of the most recently identified REDs, the pathogenesis and therapeutic intervention for CANVAS is of central significance in biomedical research. Although our understanding of CANVAS has begun to emerge, the molecular mechanism by which expanded AAGGG repeats in *RFC1* participate in pathological pathways remains largely elusive and our cognition of nucleotide repeats in the context of human diseases is still in its infancy. Besides, increasing evidence suggests that the *RFC1* AAGGG repeat expansion also associates with several other neurodegenerative diseases including the Parkinson’s disease (PD)^12, 13^, multiple system atrophy (MSA)^14–16^, late-onset ataxia (LOCA) and chronic idiopathic axonal polyneuropathy (CIAP)^17, 18^. These collectively raise an importance and urgency to tease out how the *RFC1* AAGGG repeat expansion participates in pathogenic pathways.

It has been long thought that REDs are caused by nucleotide repeats capable of forming unusual secondary structures. From the perspective of sequence context, the pathogenic *RFC1* (AAGGG)_exp_ has a higher guanine content than the non-pathogenic (AAAGG)_exp_ and (AAAAG)_exp_, and this suggests a possibility of forming G-quadruplex (G4) structure by the pathogenic (AAGGG)_exp_. G4s are secondary nucleic acid structures formed by guanine-rich nucleotides and they are increasingly recognized as key players in various biological processes associated with normal physiology and disease pathology^19, 20^. High-throughput sequencing has identified more than 700,000 G4 sites in the human genome^21^, and strikingly numbers of G4 sites are DNA repeats concatenated to neurodegenerative diseases, such as the GGGGCC repeats and CGG repeats associated with ALS/FTD and fragile X syndrome, respectively^22, 23^. It has been well established that the GGGGCC repeats form DNA and RNA G4s initiating aberrant molecular cascades during DNA replication, transcription and translation leading to ALS/FTD pathology^24–26^.

Inspired by the correlation between G4s and neurodegenerative diseases, we sought to test our hypothesis that the pathogenic *RFC1* AAGGG repeats may form G4 structures that lead to aberrant molecular cascades. In this study, we for the first time discovered and determined the high-resolution parallel G4 structure formed by the pathogenic *RFC1* AAGGG repeats. The G4 formed by short repeats tended to assemble into a higher-order structure through stacking of G-tetrads. We further show that an intramolecular G4 can be formed in a longer repetitive sequence and pose as a roadblock for polymerase processivity to cause replication stalling and transcription inhibition *in vitro*. Strikingly, a more pronounced effect on replication stalling was accompanied by longer AAGGG repeats and more stable G4 in the template. We further demonstrate that AAGGG repeats can form G4 to reduce gene expression in cells. These results point to the DNA G4 structure as a fundamental determinant of the pathogenic *RFC1* AAGGG repeats linked CANVAS and other diseases.

## Results

### The pathogenic *RFC1* AAGGG repeats form parallel G4s

To tease out how *RFC1* AAGGG repeat expansion impedes cellular function, we began with structural investigation on AAGGG repeats with various sequence lengths (Table S1). A short DNA sequence composed of two AAGGG repeats, d(AAGGG)_2_, was first examined using solution NMR and circular dichroism (CD) spectroscopy. In 20 mM K^+^, d(AAGGG)_2_ showed six guanine imino proton (G H1) signals at 11.0 to 11.6 ppm suggesting the formation of a G4 (Fig. 1b). Here we named this G4 appeared in a low K^+^ solution as G4^I^, notably which co-existed with a predominant population of random coils in 20 mM K^+^ (Fig. S1). Upon increasing the K^+^ concentration to 70 mM, the G H1 signals of G4^I^ significantly increased, but meanwhile six G H1 signals newly appeared at a more upfield region from a second type of G4 structure (G4^II^). The G4^II^ became predominant in a physiological 150 mM K^+^ concentration (Fig. 1b). CD spectra of d(AAGGG)_2_ in K^+^ solutions show positive and negative absorbance bands at 264 and 240 nm, respectively, which are signatures of parallel G4 (Fig. 1c)^20^.

The observation of six well-resolved G H1 signals in either G4^I^ or G4^II^ suggests that all the guanine residues in d(AAGGG)_2_, including G3, G4, G5, G8, G9 and G10, participated in forming G-tetrads (Fig. 1b). Yet, only six guanine residues could not form two G tetrads which is a minimum requirement of forming an intramolecular G4, and thereby both G4^I^ and G4^II^ are intermolecular G4s composed of two or more strands. We performed native polyacrylamide gel (PAGE) using two reported G4s as references, including the 12-nt TAG^27^ and 17-nt T30177^28^ that formed a three-layer dimeric G4 and a six-layer dimeric G4, respectively. Strikingly, G4^I^ and G4^II^ showed similar gel mobility to TAG and T30177, respectively (Fig. S2), suggesting that G4^I^ was dimeric whereas G4^II^ was tetrameric. This is also in line with the more upfield G H1 signals of G4^II^ owing to stacking of two G4s (Fig. 1b). The G4^II^ was likely to be formed via stacking through the 3’-terminal rather than 5’-terminal G tetrads, as swinging of A1 and A2 residues would perturb such stacking interaction.

Our attempt to determine the high-resolution structure of G4^I^ was hampered due to an unavailability of good-quality NMR spectrum for a pure G4^I^ conformation neither in a low nor high K^+^ solution (Figs. 1b and S1). It is reported that pendant groups at the termini of a G4 could disrupt staking between G tetrads and thereby prevent the formation of multimeric G4s^29^. Inspired by this, we designed two sequences via adding one and two adenine residues at the 3’-termini of d(AAGGG)_2_ to eliminate the tetrameric G4. It is worth mentioning that d(AAGGG)_2_A and d(AAGGG)_2_AA still preserves the repetitive sequence nature of the pathogenic *RFC1* AAGGG repeats. Encouragingly, d(AAGGG)_2_AA formed a pure dimeric G4 exhibiting similar NMR signal features to the G4^I^ of d(AAGGG)_2_ in 150 mM K^+^ (Figs. 1d and S3), and the CD spectra indicate a parallel G4 of d(AAGGG)_2_AA (Figs. 1e and S4).

### NMR solution structure determination of the *RFC1* G4

To provide further insights into the conformation and self-assembly property of the *RFC1* G4, we determined the solution NMR structures of d(AAGGG)_2_AA. Two-dimensional NMR spectra, including nuclear Overhauser effect spectroscopy (NOESY), double quantum-filtered correlation spectroscopy (DQF-COSY), total correlation spectroscopy (TOCSY) and heteronuclear multiple bond correlation (HMBC), were acquired using unlabeled d(AAGGG)_2_AA to assign H8 and H1’ (Fig. 2a), sugar protons and adenine H2. Besides, six sequences with a 6% ^15^N-isotope labeling at the designated position of G3, G4, G5, G8, G9 and G10, respectively, were prepared, and the guanine H1 resonances were assigned via ^1^H-^15^N heteronuclear single quantum coherence (HSQC) spectra (Fig. 2b).

**Fig. 2.**
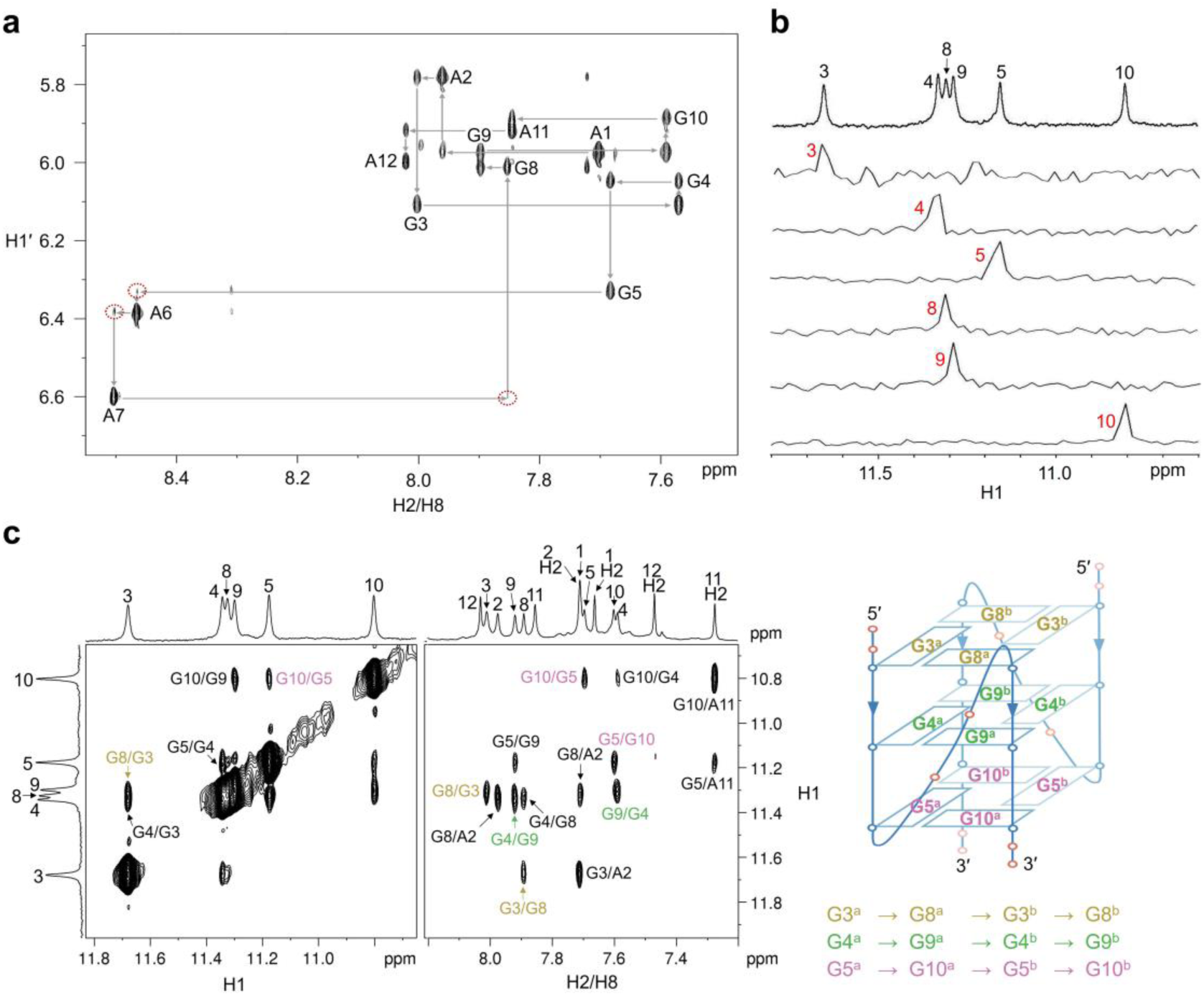
NMR spectra of d(AAGGG)_2_AA. **a** H6/H8-H1’ fingerprint region from the NOESY spectrum. **b** ^15^N-filtered HSQC spectra of site-specific 6% ^15^N-labeled d(AAGGG)_2_AA. **c** H1-H1 and H8-H1 regions from the NOESY spectrum show NOEs connections in each G tetrad.

The solution NMR structures of d(AAGGG)_2_AA were calculated using restrained molecular dynamic (rMD) simulations based on 252 NOE-derived proton-proton distance, 48 hydrogen bond, 4 dihedral angle, 24 torsion angle, 36 G-tetrad planarity and 72 chirality restraints. Ten structures with the lowest total energies were selected as the final representative ensemble (Fig. 3a), and they were well-converged with a heavy atom root-mean-squire-deviation (RMSD) of 0.5 ± 0.1 and 1.0 ± 0.2 Å for the G-tetrad core and all residues, respectively (Table 1).

**Fig. 3.**
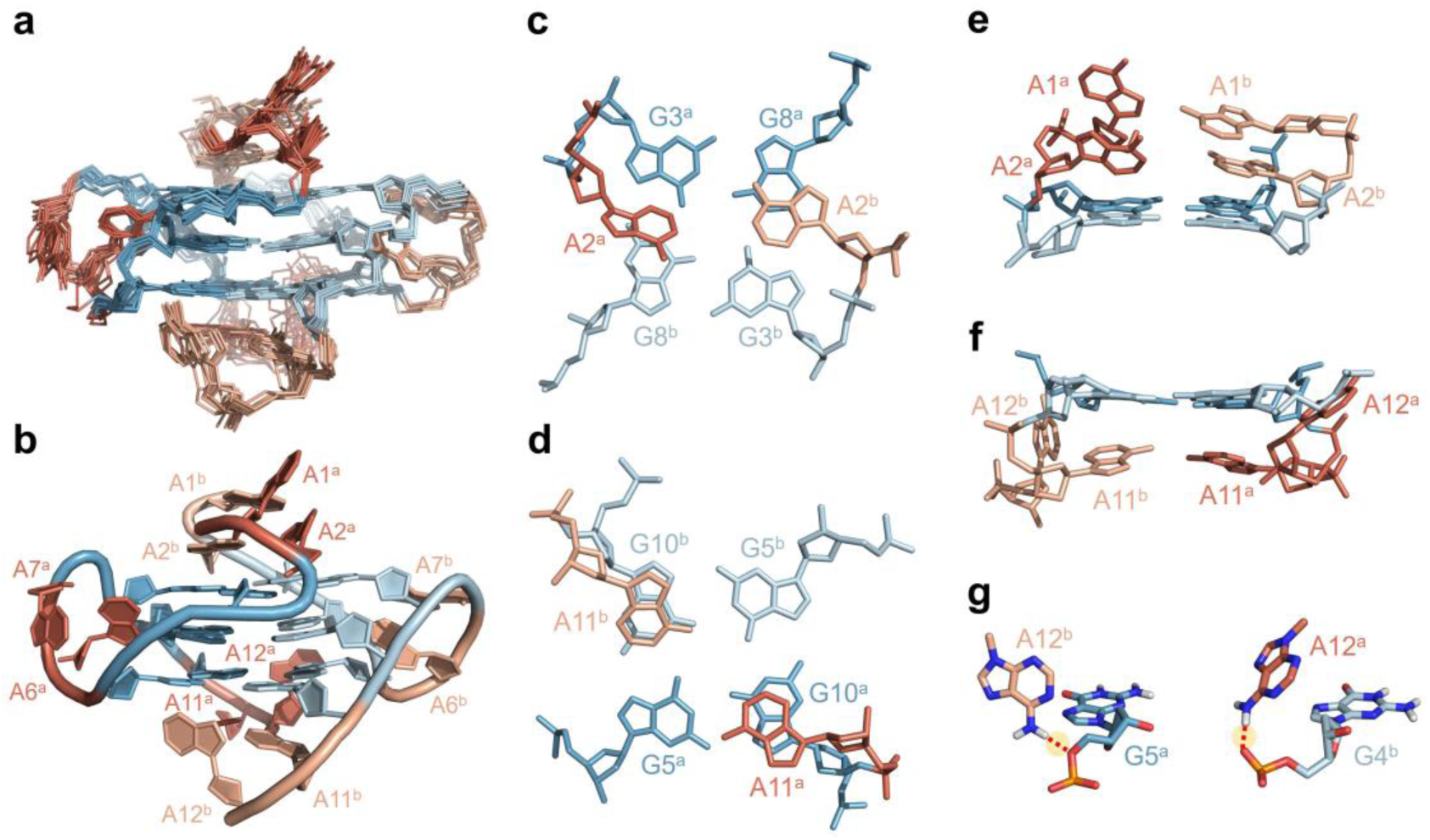
Solution NMR structures of the G4 formed by d(AAGGG)_2_AA. **a-b** Superimposed 10 representative structures and a single structure with the lowest total energy. **c-f** Conformations of the 5’-terminal and 3’-terminal adenine residues. **g** Hydrogen bonds of A12^b^ H61···G5^a^ O5′ and A12^a^ H61···G4^b^ OP2.

**Table 1.** NMR restraints and refinement statistics of the G4 of d(AAGGG)_2_AA.

|  |  |  |
| --- | --- | --- |
| **Structural restraints** |  |  |
| NOE-derived distance |  | 252 |
| Glycosidic torsion angle ($\chi$) | | 24 |
| Sugar torsion angle (H1'-C1'-C2'-H2') |  | 4 |
| Hydrogen bond |  | 48 |
| Planarity |  | 36 |
| Chirality |  | 72 |
| **Restraint satisfaction** |  |  |
| Number of distance violations > 0.2 Å |  | 0 |
| Maximum distance violation (Å) |  | 0.192 |
| Number of angle violations > 5° |  | 1 |
| Maximum angle violation (°) |  | 5.4 |
| **Deviations from ideal covalent geometry** |  |  |
| Bonds (Å) |  | 0.0079 ± 0.0004 |
| Angles (°) |  | 2.4 ± 0.03 |
| **Pairwise heavy atom RMSD (Å)** |  |  |
| All residues |  | 1.0 ± 0.2 |
| G-tetrad core |  | 0.5 ± 0.1 |

The d(AAGGG)_2_AA formed a dimeric parallel G4 containing three G-tetrad layers (Fig. 3b), including G3^a^→G8^a^→G3^b^→G8^b^, G4^a^→G9^a^→G4^b^→G9^b^ and G5^a^→G110^a^→G5^b^→G10^b^ wherein the superscript a and b represent two respective DNA chains, and the arrow indicates the direction of Hoogsteen H-bond donor-to-acceptor in each G-tetrad supported by internucleotide G H1-H8 NOE connections (Figs. 2c). To connect the three G-tetrads, the phosphodiester backbones of A6^a/b^ and A7^a/b^ are highly twisted, which is consistent with the abnormally weak NOEs of G5 H1′-A6 H8, A6 H1′-A7 H8 and A7 H1′-G8 H8 (Fig. 2a). More intriguingly, the 5’-termi and 3’-termi exhibit different conformational plasticity. At the 5’-termi, A2^a^ and A2^b^ stacked on the G3^a^·G8^a^·G3^b^·G8^b^ tetrad, whereas A1^a^ and A1^b^ further stacked on A2^a^ and A2^b^, respectively, with A1^a^ and A2^a^ being nearly perpendicular to the G-tetrad (Fig. 3c, e). These are also supported by NOEs of A2 H2-G3/G8 H1 (Fig. 2c) and A1 H2/H8-A2 H8 (Fig. S5). At the 3’-termini, A11^a^ and A11^b^ stacked on the G5^a^·G10^a^·G5^b^·G10^b^ tetrad (Fig. 3d, f) as agreed with NOEs of A11 H2-G5/G10 H1 (Fig. 2c), while A12^a^ and A12^b^ instead of stacking on A11^a^ and A11^b^, folded into the minor groove to form hydrogen bonds with G4^b^ and G5^a^ via A12^b^ H61···G5^a^ O5′ and A12^a^ H61···G4^b^ OP2, respectively (Fig. 3f, g). The distinctive conformational plasticity between 5’ and 3’-termini of d(AAGGG)_2_AA is in consistence with a recent finding that the 3’-terminal residues display more dynamics than 5’-terminal residues in parallel G4s^30^. This structural feature fitly rationalizes why addition of only one adenine residue at the 3’-termini to d(AAGGG)_2_ could not prevent the formation of a higher-order tetrameric G4 (Fig. S3), as A11^a^ and A11^b^ formed a blunt end allowing stacking between two G4s (Fig. 3d, f).

### N-methyl mesoporphyrin IX (NMM) binds and stabilizes the G4 formed by pathogenic *RFC1* AAGGG repeats

Prior to exploring the functional consequence of the pathogenic *RFC1* AAGGG repeats, we sought to seek for a small-molecule ligand that can bind and stabilize the G4 formed by AAGGG repeats. Three typical G4 ligands^31^, including pyridostatin (PDS), TMPyP4 and NMM, were tested using CD and fluorescence spectroscopy. It was found that addition of PDS and TMPyP4 to d(AAGGG)_2_AA weakened the CD absorbance band at 264 nm (Fig. 4a), suggesting that they could bind to d(AAGGG)_2_AA but alter the parallel G4 topology. Addition of NMM did not perturb the G4 structure of d(AAGGG)_2_AA (Fig. 4a). Binding to a G4 brings about π-π conjugation effect that enhances the fluorescence of NMM at an emission wavelength at 610 nm^32^. Titration of d(AAGGG)_2_AA to NMM drastically enhanced the fluorescence intensity at 610 nm, consolidating the binding between NMM and d(AAGGG)_2_AA G4 (Fig. 4b). Further, CD melting experiment reveals a drastic increase in the melting temperature (*T_m_*) of d(AAGGG)_2_AA G4 by ∼11.5 °C in the presence of excessive NMM (Figs. 4c, S6 and Table S2).

**Fig. 4.**
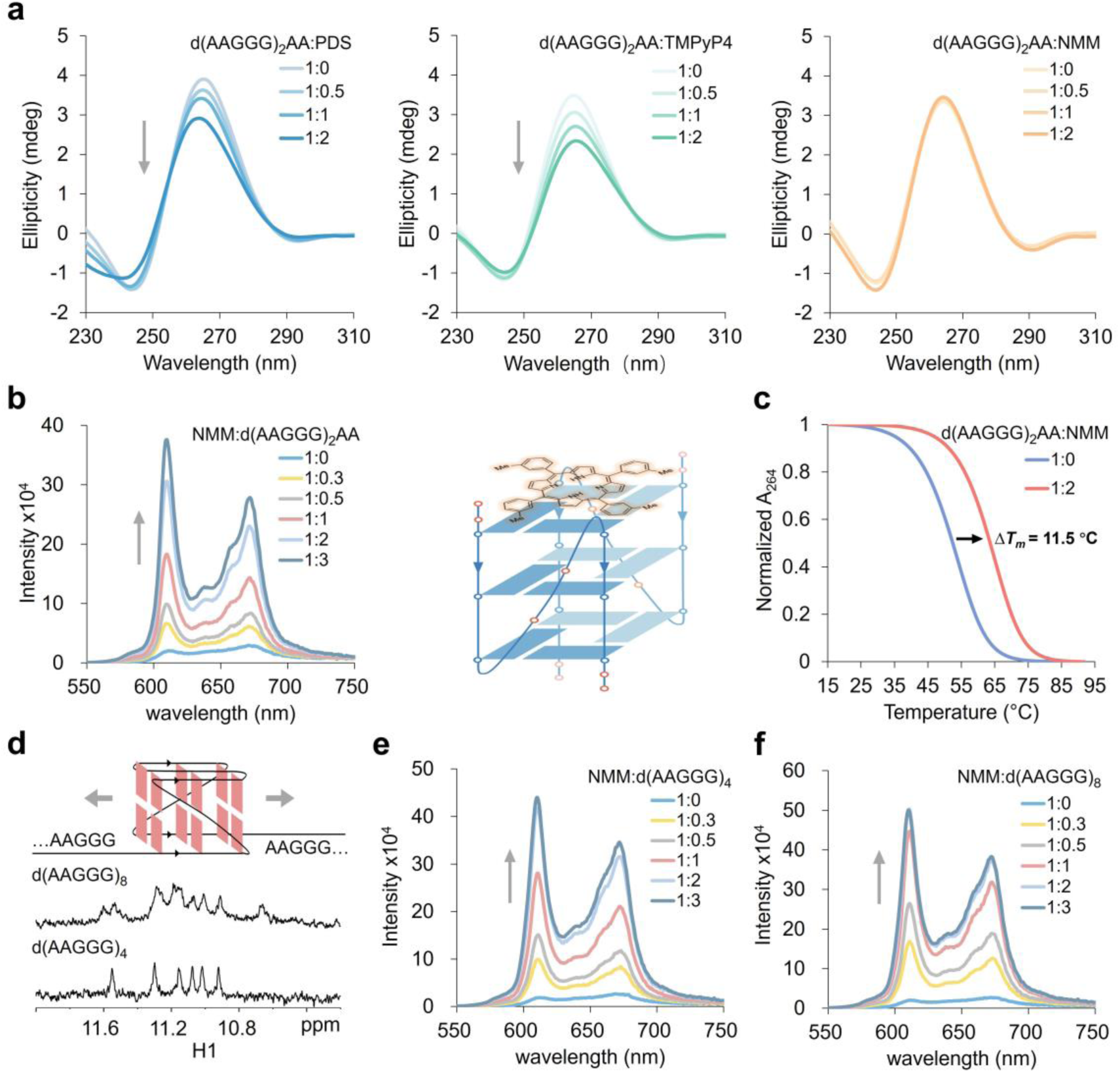
NMM binds and stabilizes the DNA G4 formed by AAGGG repeats. **a** CD spectra of ligand titration to d(AAGGG)_2_AA. **b** Fluorescence spectra NMM upon d(AAGGG)_2_AA titration. **c** CD melting curves of d(AAGGG)_2_AA with and without NMM. **d** 1D ^1^H NMR spectra of d(AAGGG)_4_ and d(AAGGG)_8_. **e-f** Fluorescence spectra of NMM upon d(AAGGG)_4_ and d(AAGGG)_8_ titration.

We also examined if longer AAGGG repeats could form an intramolecular parallel G4. In 150 mM K^+^, d(AAGGG)_4_ and d(AAGGG)_8_ were found to form parallel G4s as collectively suggested by their G H1 signals at ∼10.5 to 11.5 ppm (Fig. 4d) and CD spectral features (Fig. S7). However, we noticed that d(AAGGG)_4_ still exhibited six G H1 signals which are similar to those of d(AAGGG)_2_AA (Figs. 1b, 4d), and the native PAGE showed that d(AAGGG)_4_ indeed formed a dimeric G4 (Fig. S8). Strikingly, the longer d(AAGGG)_8_ formed an intramolecular three-layer G4 as suggested by twelve G H1 signals (Fig. 4d). The binding of NMM to d(AAGGG)_4_ and d(AAGGG)_8_ was further evidenced by enhanced fluorescence at 610 nm upon DNA titration (Fig. 4e, f). In short, obtaining the G4-stablizing ligand NMM and consolidating the formation of G4 in longer AAGGG repeats lay a foundation for us to examine the functional consequence of the pathogenic *RFC1* AAGGG repeats.

### Formation of G4 in *RFC1* AAGGG repeats impeded replication and transcription *in vitro*

Numbers of neurodegenerative diseases leading to ataxia and neuropathy are linked to DNA damage and abnormal repair pathways^33, 34^. Non-B DNA structures, in particular the thermodynamically stable G4s, can pose obstacles for polymerase processivity during DNA replication, thus increasing polymerase stalling and replication errors^35, 36^ We therefore examine if the pathogenic *RFC1* AAGGG repeats can form G4 in an elongated template to impede DNA polymerase processivity. First, we established an *in vitro* replicational assay by placing AAGGG repeats with variable lengths, (AAGGG)_4_ and (AAGGG)_8_, downstream of the priming site (Fig. 5a). Two reference templates containing (AAAAG)_4_ and (AAAAG)_8_ were also prepared. Remarkably, (AAAAG)_4_ or (AAAAG)_8_ is a part of non-pathogenic sequence in normal *RFC1* alleles (Fig. 1a) and they are supposed to be uncapable of forming G4 structure. The 5’-Cy5 labeled primer was extended by Klenow fragment (KF) of DNA polymerase I with 5′ to 3′ polymerase and 3′ to 5′ exonuclease activities, and this KF has been used to study effects of various types of non-B DNAs on polymerase processivity *in vitro*^37, 38^.

**Fig. 5.**
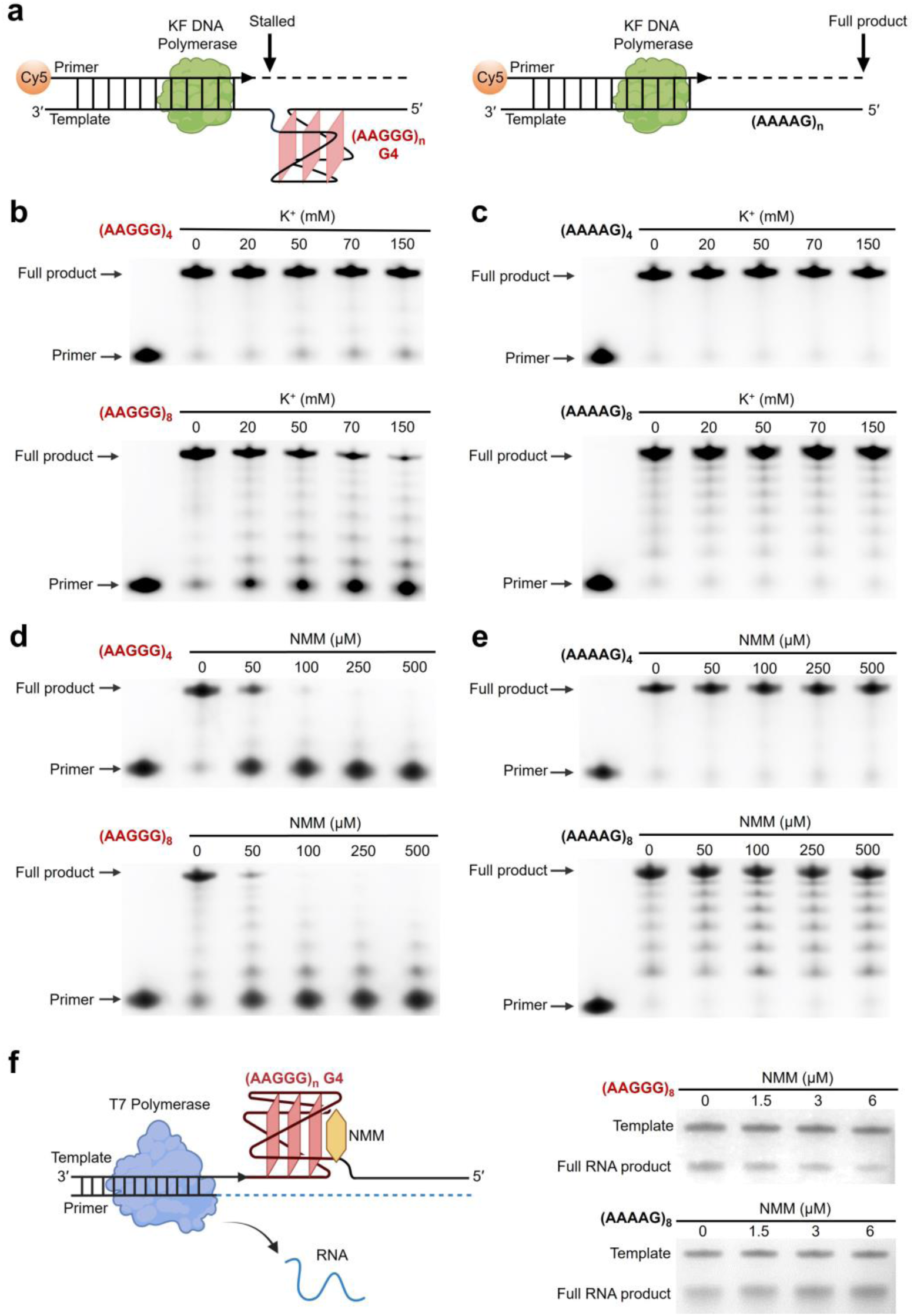
Formation of DNA G4 in *RFC1* AAGGG repeats impeded polymerase processivity during replication and transcription. **a** Schematic of the *in vitro* KF extension assay on primer-template models containing pathogenic AAGGG repeats and non-pathogenic AAAAG repeats. **b-e** Denaturing gels show KF extension products under various concentrations of K^+^ and NMM. **f** *In vitro* T7 transcription assay on primer-template models containing (AAGGG)_8_ and (AAAAG)_8_ under various concentrations of NMM.

The *in vitro* replicational products were resolved by denaturing PAGE. For the template containing (AAGGG)_4_, increasing K^+^ had a slight impeding effect on the polymerase processivity and few nonextended products were observed in 50 to 150 mM K^+^ (Fig 5b). Strikingly, the template containing a longer (AAGGG)_8_ showed significantly reduced full-length extension products in 70 to 150 mM K^+^ (Fig 5b), suggesting that K^+^-induced G4 formation severely impaired polymerase processivity. It also appears that the replication stalling is repeat-length-dependent under a physiologically relevant ionic condition, and this trend points to a possible mechanism for the fact that disease manifests only when the number of expanded repeats exceeds a certain threshold. In contrast, the polymerase processivity was not affected by K^+^ concentration for the reference templates containing non-pathogenic AAAAG repeats, and the full-length extension products were efficiently synthesized (Fig. 5c). We noted that the reference template containing (AAAAG)_8_ also generated few truncated extension products, and this may be attributed to an intrinsic slowing down of DNA polymerase processivity when encountering a repetitive sequence^39^.

As it has been demonstrated that the NMM could bind and substantially stabilize the G4 formed by AAGGG repeats (Fig. 4), we also performed the *in vitro* replicational assay under various concentrations of NMM to examine the effect of G4 stabilization on replication stalling. The impeding effect of NMM-induced G4 stabilization was tremendous for both the templates containing (AAGGG)_4_ and (AAGGG)_8_, and no appreciable full-length extension product was observed under 100 μM NMM (Fig. 5d). The polymerase progressivity was not affected by non-pathogenic (AAAAG)_4_ and (AAAAG)_8_ under various concentrations of NMM (Fig. 5e).

We also established an *in vitro* transcriptional assay by placing (AAGGG)_8_ downstream of a T7 promoter as the transcription template. It showed that the NMM-induced G4 stabilization leads to a reduction in the full-length RNA product (Fig. 5f). As expected, the transcription was not inhibited by NMM for the non-pathogenic (AAAAG)_8_. Collectively, the *in vitro* replicational and transcriptional assays demonstrated that the pathogenic *RFC1* AAGGG repeats can form G4 structure in an elongated template to impede polymerase processivity. Moreover, the impeding effect became more pronounced when the number of AAGGG repeats was increased and the thermodynamic stability of G4 was enhanced.

### Formation of G4 in *RFC1* PRE reduces gene expression in cells

Two latest works have reported reduced RFC1 protein levels in CANVAS patients yet the underlying molecular mechanism remains elusive^10, 11^. As the G4 formation in AAGGG repeats impairs transcription *in vitro* (Fig. 5f), we wondered if the reduced protein level is also related to G4 formation. To further understand this, we constructed a live-cell gene expression reporter. A pBudCE4.1 vector-based plasmid construct expresses EGFP under the CMV promoter to serve as a gene expression reporter, and expresses nuclear localization signal-mTagBFP2 under the EF-1α promoter to serve as an internal reference. The pathogenic *RFC1* (AAGGG)_8_ or non-pathogenic (AAAAG)_8_ was inserted upstream of the EGFP sequence (Fig. 6a).

**Fig. 6.**
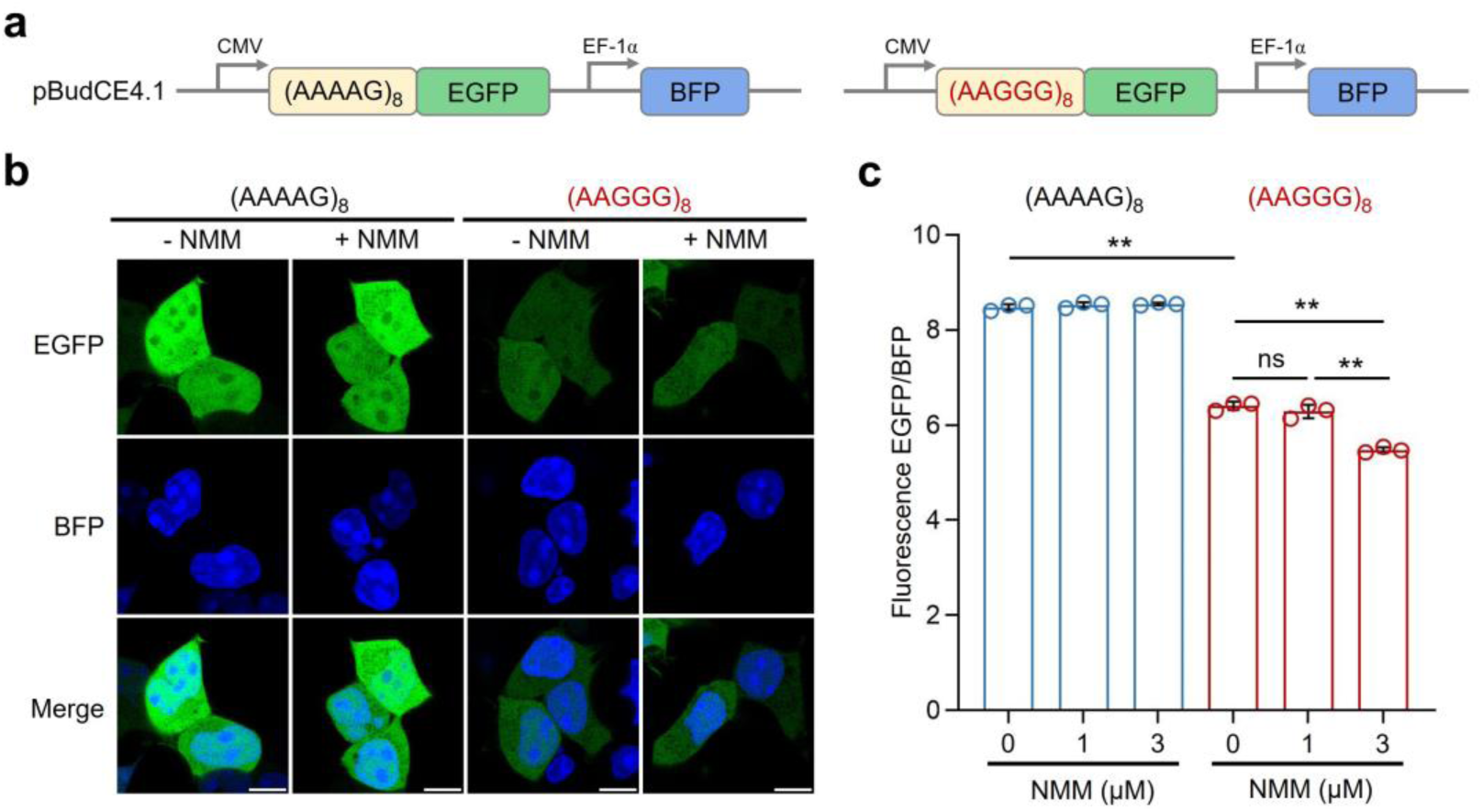
Formation of G4 in *RFC1* AAGGG repeats reduced gene expression in cells. **a** Schematic of the live cell-based gene expression reporter. **b-c** Confocal microscopy (scale bars, 10 µm) and flow cytometry assays on HEK293T cells expressing (AAAAG)_8_ and (AAGGG)_8_ constructs with NMM treatment. **P < 0.01.

After 4 h post-transfection with the plasmid construct, HEK293T cells were incubated in a culture medium containing 0 or 3 μM NMM for 24 h. Using the BFP fluorescence as an internal reference under confocal microscopy, cells transfected with the reference (AAAAG)_8_ construct did not show obvious difference in EGFP fluorescence regardless of NMM treatment (Fig. 6b). In contrast, the EGFP fluorescence became much weaker in cells transfected with the (AAGGG)_8_ construct, suggesting a reduced level of EGFP expression. To further verify if it was the G4 formation in AAGGG repeats contributed to the weakened EGFP fluorescence, we quantified EGFP expression by flow cytometry based on the relative fluorescence intensity of EGFP/BFP. For cells transfected with the reference (AAAAG)_8_ construct, there was no significant difference in EGFP expression between groups treated with 0, 1 and 3 µM NMM (Fig. 6c). However, cells transfected with the (AAGGG)_8_ construct even without NMM treatment showed a significant reduction in EGFP expression by ∼25% comparing to cells transfected with the (AAAAG)_8_ construct. This suggests that AAGGG repeats may naturally form G4 structure in cells to exert functional consequence. More strikingly, treatment with NMM (3 µM) to cells expressing the (AAGGG)_8_ construct further lowered the EGFP level by ∼14% (Fig. 6c), further confirming that the G4 formation indeed contributed to the reduced gene expression.

## Discussion

Since the latest discovery of pathogenic *RFC1* AAGG repeat expansion linked to CANVAS^4–7^ and several other neurodegenerative diseases including the PD^12, 13^ and MSA^14–16^, few progress has been achieved towards a clear understanding on molecular mechanisms of expanded AAGGG repeats in disease pathology. The biological consequence of nucleic acids is not only determined by their primary sequence, but also the structural property. Structural diversity of nucleic acids governs the fate of various biological processes concatenated to physiological function and pathology^23, 40^. Here we find that formation of DNA G4 structure in the pathogenic *RFC1* AAGGG repeats underlies several aberrant molecular cascades including replication stalling, transcription inhibition and gene expression (Fig. 7).

**Fig. 7.**
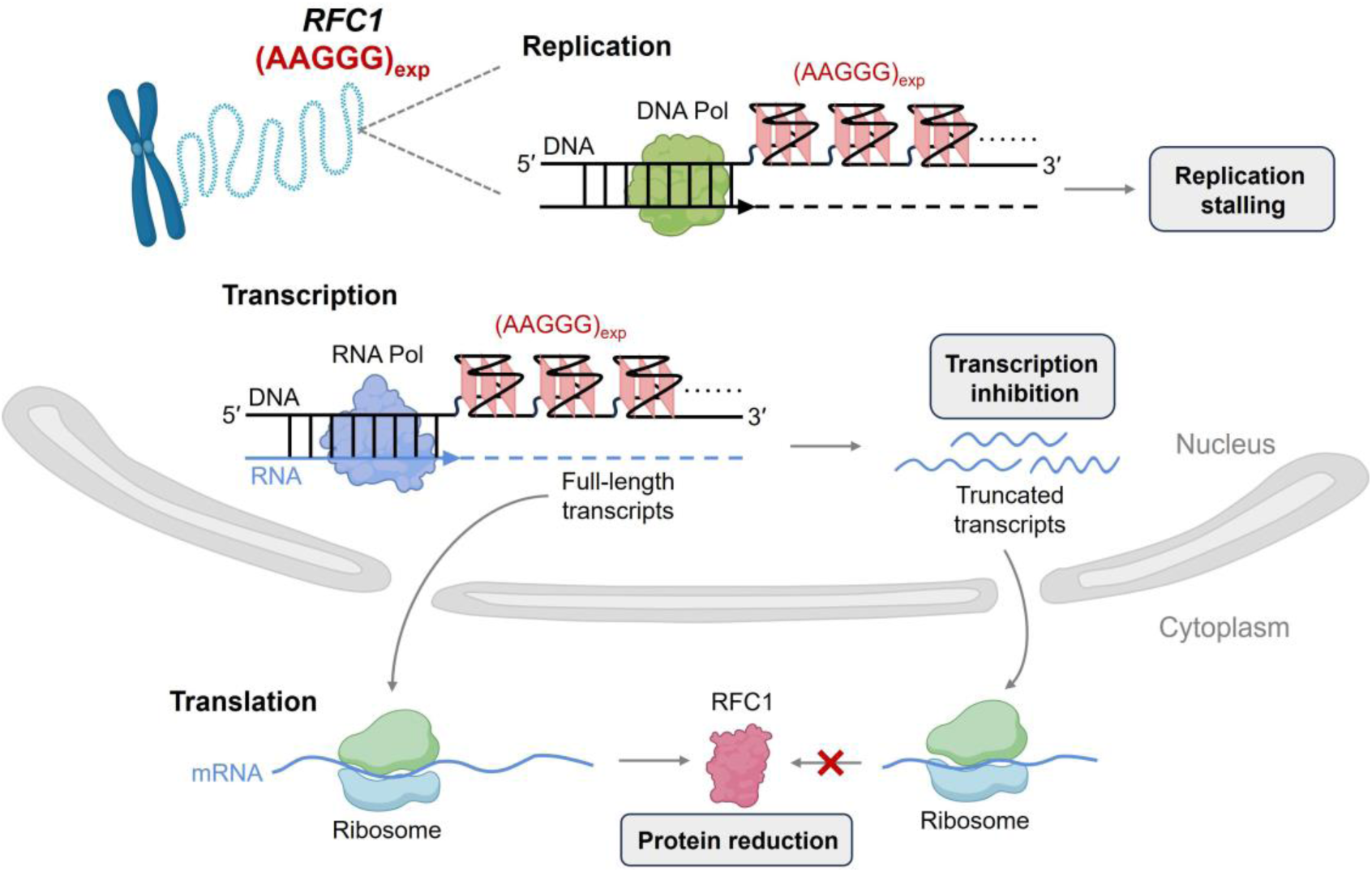
A proposed model for the functional consequence of G4 formation in the pathogenic *RFC1* AAGGG repeats. The expanded AAGGG repeats form DNA G4 structure(s) that impede polymerase processivity, causing replication stalling and transcription inhibition, which further partially contributes to reduced RFC1 protein levels.

In the mutated *RFC1* alleles of CANVS patients, the number of pathogenic AAGGG repeats ranges from hundreds to thousands^4–7^. Although a correlation between the repeat length and clinical onset or progression has yet to be established, it is a common feature that all REDs identified so far manifest only beyond a critical number of repeats^1^. Our *in vitro* replicational assay shows that AAGGG repeats in the template could impede polymerase processivity and cause replication stalling in a repeat-length-dependent manner (Fig. 5). It is not difficult to understood that longer repeats have a higher possibility to form G4 with an increasing number of G tracts. Besides, we found that the G4s formed by AAGGG repeats tended to self-assembly (Figs. 1, S2, S3, S8), therefore it is possible that longer repeats will allow the formation of adjacent G4s posing as a more complex roadblock for polymerase processivity (Fig. 7). The mutated *RFC1* allele harboring hundreds to thousands of AAGGG repeats, which potentially form multiple G4 structural obstacles, is anticipated to suffer from more pronounced replication errors and DNA damage *in vivo*.

Our results showed transcription inhibition *in vitro* and reduced gene expression in cells due to formation of G4 in AAGGG repeats (Figs. 5f, 6). These collectively pointed to a new role of DNA G4 within the mutated *RFC1* PRE in impairing protein production, and could provide a plausible mechanism for the recently reported RFC1 protein reduction in CANVAS patients^10, 11^. With a more prudent consideration, the reduced protein levels detected in our live-cell gene expression reporter may also be partially contributed by the formation RNA G4 in mRNA transcripts harboring AAGGG repeats, as the downregulation of gene expression by RNA G4 is not uncommon^41–43^. In REDs, the RNA transcripts containing expanded repeats lead to cellular dysfunction through more complex mechanisms, such as sequestration of RNA-binding proteins and formation of nuclear foci^44, 45^. There is a possibility that the RNA G4 structure of *RFC1* PRE may also elicit detrimental effects on cellular function and it undoubtedly worths for further investigations.

Apart from providing a molecular mechanism by which the pathogenic *RFC1* PRE participates in several aberrant molecular cascades, more importantly for the first time we have resolved the high-resolution structures of the G4 formed by pathogenic AAGGG repeats. An intriguing structural feature of the G4 is a higher conformational plasticity of the 3’-terminal residues than 5’-terminal residues, thus two G4s tended to assemble into a higher-order G4 structure through stacking at 3’-termini. The DNA G4 formed in *RFC1* PRE appears to elicit detrimental effects on several key biological processes, and thus may be a potential therapeutic target. It has been recently reported that proteins or helicases, such as the Cockayne Syndrome B protein and DEAH-Box helicase 9, could resolve pathogenic DNA G4 structures to restore normal cellular function^46, 47^. The unprecedented high-resolution structures of G4 formed by AAGGG repeats will facilitate the discovery of helicase and small-molecule ligands to resolve the culpable G4 structure in *RFC1* PRE for therapeutic intervention.

## Methods

### Sample preparation

DNA oligonucleotides were purchased from Sangon Biotech Co. Ltd. (Shanghai, China) with HPLC purification, and further purified in our laboratory using diethylaminoethyl sephacel anion exchange column and centrifugal desalting. The 6% site-specific ^13^C/^15^N isotopically labelled oligonucleotide were synthesized on K&A H8 synthesizer using the 2’-deoxyadenosine phosphoramidite (98% ^15^N) purchased from Cambridge Isotope Laboratories (USA), and purified by denaturing PAGE, electro-elution, diethylaminoethyl sephacel anion exchange column, and centrifugal desalting. DNA samples were quantified using a NanoDrop microvolume spectrophotometer.

### NMR spectroscopy

NMR samples contained 100 µM DNA (1D experiments) or 800 µM DNA (2D experiments), 1 mM NaPi (pH 7), 0 to 150 mM KCl, and 90% H_2_O/10% D_2_O or 99.96% D_2_O. NMR spectra were acquired on a Bruker AVANCE 600 MHz spectrometer and analyzed using TopSpin software. Excitation sculpting and pre-saturation water suppression methods were applied for 90% H_2_O/10% D_2_O and 99.96% D_2_O samples, respectively. 2D NMR experiments, including NOESY (mixing times of 100, 200 and 300 ms), DQF-COSY, TOCSY (mixing time of 75 ms), ^1^H-^13^C HSQC (^1^*J*_C,H_ of 180 Hz), and ^1^H-^15^N HSQC (^1^*J*_N,H_ of 90 Hz) were acquired at 25 °C. The H8 and H1’ resonances were assigned from NOESY, and the H2’, H2’’, H3’, H4’, H5’ and H5’’ resonances were assigned from DQF-COSY and/or TOCSY spectra. The guanine H1 resonances were unambiguously assigned from ^1^H-^15^N HSQC spectra using the site-specific 6% ^15^N-labeled DNA samples.

### Structural calculation

NOE-derived distance restraints of d(AAGGG)_2_AA were obtained based on the NOE cross-peaks integrated from NOESY spectra (mixing time of 100-300 ms) as described previously. Glycosidic torsion angles of 90-330° for *anti* and -90-90° for *syn* were applied based on intranucleotide H8–H1’ NOE intensity and C8 chemical shifts. H1’-C1’-C2’-H2’dihedral angles were determined by ^3^*J*_H1’-H2’_ coupling constants measured from the DQF-COSY spectrum and Karplus equation^48^. The structural calculations were carried out on AMBER18^49^ using the bsc1 force field^50^. The starting models were energy minimized, and then subjected to restrained molecular dynamic (rMD) simulations and restrained energy minimization (rEM). During the 35-ps annealing process, the system temperature increased from 300 to 600 K from the first 5 ps, maintained at 600 K for 20 ps, decreased to 300 K within 5 ps, and stayed at 300 K for 5 ps. The structural coordinates were then subjected to rEM by 200 steps of the steepest descent and conjugated gradient minimization steps until the energy gradient difference between successive minimization steps was smaller than 0.1 kcal·mol^-1^ ·Å^-2^. Among the 500 structures calculated with random seeds, 10 structures with the lowest total energies were selected as the final representative ensemble. RMSD values were calculated using the *suppose* module of AMBER.

### Native PAGE

Native PAGE was performed using 20% polyacrylamide gels supplemented with 1× TBE buffer at room temperature. DNA loading samples contained 0.1 mM DNA in 1 mM NaPi (pH 7) and variable concentrations of K^+^ as stated in the figure legends. DNA bands were visualized by staining the gels with stains-all solutions.

### CD spectroscopy

CD samples were prepared to contain 20 μM DNA in 1 mM NaPi (pH 7) and variable K^+^ concentrations as stated in the figure legends. CD spectra were recorded on a Chirascan V100 spectropolarimeter using 1 mm path length quartz cuvette and 1 nm bandwidth at room temperature. The blank correction was made by subtracting the buffer spectrum. For CD melting experiments, the CD ellipticity at 264 nm was recorded from 15 to 95 °C with a heating rate of 1 °C/min. Thermodynamic parameters were determined by fitting the CD melting curves using a two-state transition model^51^.

### Fluorescence experiments

Fluorescence experiments were performed on an Edinburgh instruments FLS1000 spectrometer at room temperature. The stock DNA sample of d(AAGGG)_2_AA G4 was prepared in 1 mM NaPi (pH 7) and 150 mM KCl. The stock DNA solution was titrated into the ligand solution (1 µM PDS, TMPyP4 or NMM) in 1 mM NaPi (pH 7) and 150 mM KCl. The complex solution was mixed well and equilibrated for 5 min. The emission spectra were measured using a 10 mm path length cuvette with an excitation wavelength of 393 nm and a recorded range of 550-750 nm.

### *In vitro* replicational assay

The mixture of template and 5’-Cy5 labeled primer (1:1 equivalent) were annealed by heating at 95 °C for 5 min and cooling to room temperate for overnight. For studying the effect of NMM-stabilized G4 formation on polymerase processivity, NMM was added into the reaction mixture at various concentrations and incubated at room temperature for 2 h. Primer extension was performed for 1 h at 37 °C in a 20 µL reaction buffer containing 50 µM primer-template, 1.25 mM dNTPs, 0.2 U/µL KF polymerase, 5 mM NaCl, 1 mM Tris-HCl, 50 mM KCl and 1 mM MgCl_2_. For studying the effect of K^+^-stabilized G4 formation on polymerase processivity, the reaction buffer contained 0 to 150 mM K^+^. The primer extension products were resolved using 10% denaturing PAGE and visualized by Cytiva Amersham ImageQuant 800.

### *In vitro* transcriptional assay

The mixture of primer and template was prepared and treated with NMM as described above. Transcription was performed for 14 h at 37 °C in a 20 µL reaction buffer containing 3 µM primer-template, 5 mM Tris-HCl pH 8, 15 mM NaCl, 10 mM MgCl_2_, 10 mM rNTPs, 9 U/µL T7 RNA polymerase, 50 mM KCl and 0/1.5/3/6 µM NMM. The transcripts were resolved using 10% denaturing PAGE and visualized by staining the gel with stains-all solution.

### Cell culture and transfection

HEK293T cells were cultured in DMEM supplemented with 10% (v/v) fetal bovine serum (FBS) at 37 °C with 5% CO_2_. Transient transfection of the pBudCE4.1 plasmid construct into HEK293T cells were conducted using Lipofectamine 3000 reagent (Thermo Fisher Scientific) according to the manufacturer’s recommendations. In brief, 1000 ng plasmid, 1 μL P3000 in 25 μL Opti-MEM (Gibco), and 1 μL Lipofectamine 3000 in 25 μL Opti-MEM were mixed and incubated for 15 min at room temperature. After culturing for 24 h, cells were harvested for confocal microscopy and flow cytometry analyses.

### Confocal microscopy

HEK293T cells were plated in eight-well microscope slide with 2×10^5^ cells/well and cultured for overnight. After 4 h post-transfection with plasmid, the cells were cultured for 24 h in a fresh medium containing 0/3 µM NMM with 0.1% DMSO. Then the cells were washed with PBS. The fresh Phenol Red-free DMEM media with 10% FBS was added prior to confocal microscopic experiment (Leica Stellaris 8). A 405 nm laser excitation was used to image BFP and a 488 nm laser excitation was used to image EGFP.

### Flow cytometry

HEK293T cells were plated in a six-well plate with 5×10^5^ cells/well and cultured for overnight. After 4 h post-transfection with plasmid, the cells were cultured for 24 h in a fresh medium containing 0/1/3 µM NMM with 0.1% DMSO. Then the cells were washed with PBS and resuspended in fresh PBS with 4% FBS for flow cytometry experiment (Thermo Fisher Attune NxT).

### Statistical Analysis

Statistical analyses were performed using the GraphPad Prism 5.0. Data are given as means ± SD by three replicative experiments. Quantitative analysis was performed using two-tailed Student’s t test. P value less than 0.01 was taken as statistically significant.

## Supplementary materials for

**Table S1.**
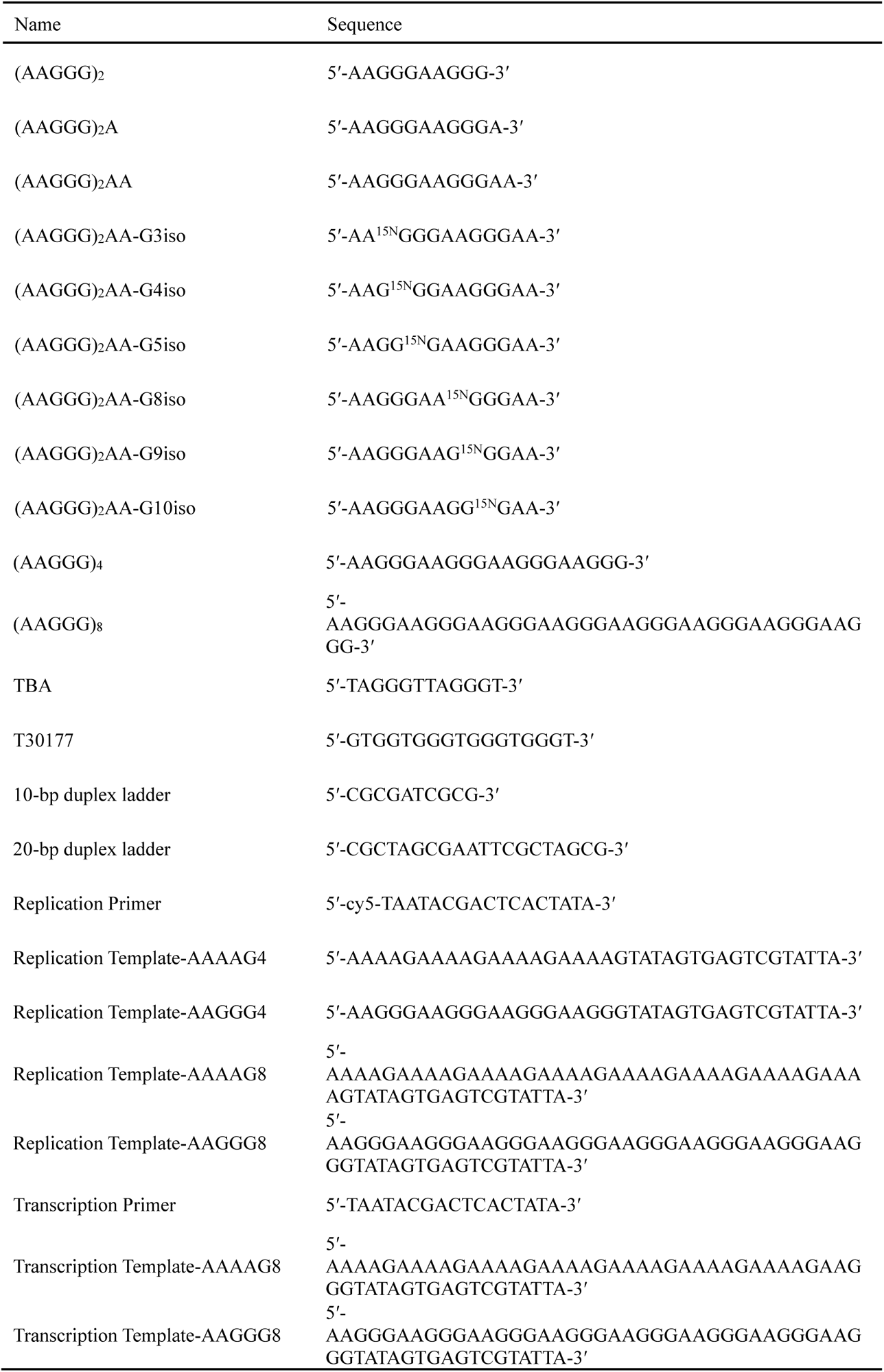
DNA sequences used in this study.

**Table S2.** Thermodynamic parameters of d(AAGGG)_2_AA G4 in the presence of NMM.

| DNA:NMM | $T_m$ (°C) | $\Delta H^\circ$<br>(kcal·mol<sup>-1</sup>) | $\Delta S^\circ$<br>(cal·K<sup>-1</sup>·mol<sup>-1</sup>) | $\Delta G^\circ_{298K}$<br>(kcal·mol<sup>-1</sup>) |
| --- | --- | --- | --- | --- |
| 1:0 | 52.3 ± 0.1 | -59 ± 3 | -182 ± 10 | -5.0 ± 0.3 |
| 1:0.5 | 58.2 ± 0.5 | -54 ± 8 | -163 ± 24 | -5.4 ± 0.9 |
| 1:1 | 61.8 ± 0.3 | -62 ± 4 | -186 ± 12 | -6.9 ± 0.4 |
| 1:2 | 63.76 ± 0.08 | -69 ± 3 | -204 ± 10 | -7.9 ± 0.4 |

**Fig. S1.**
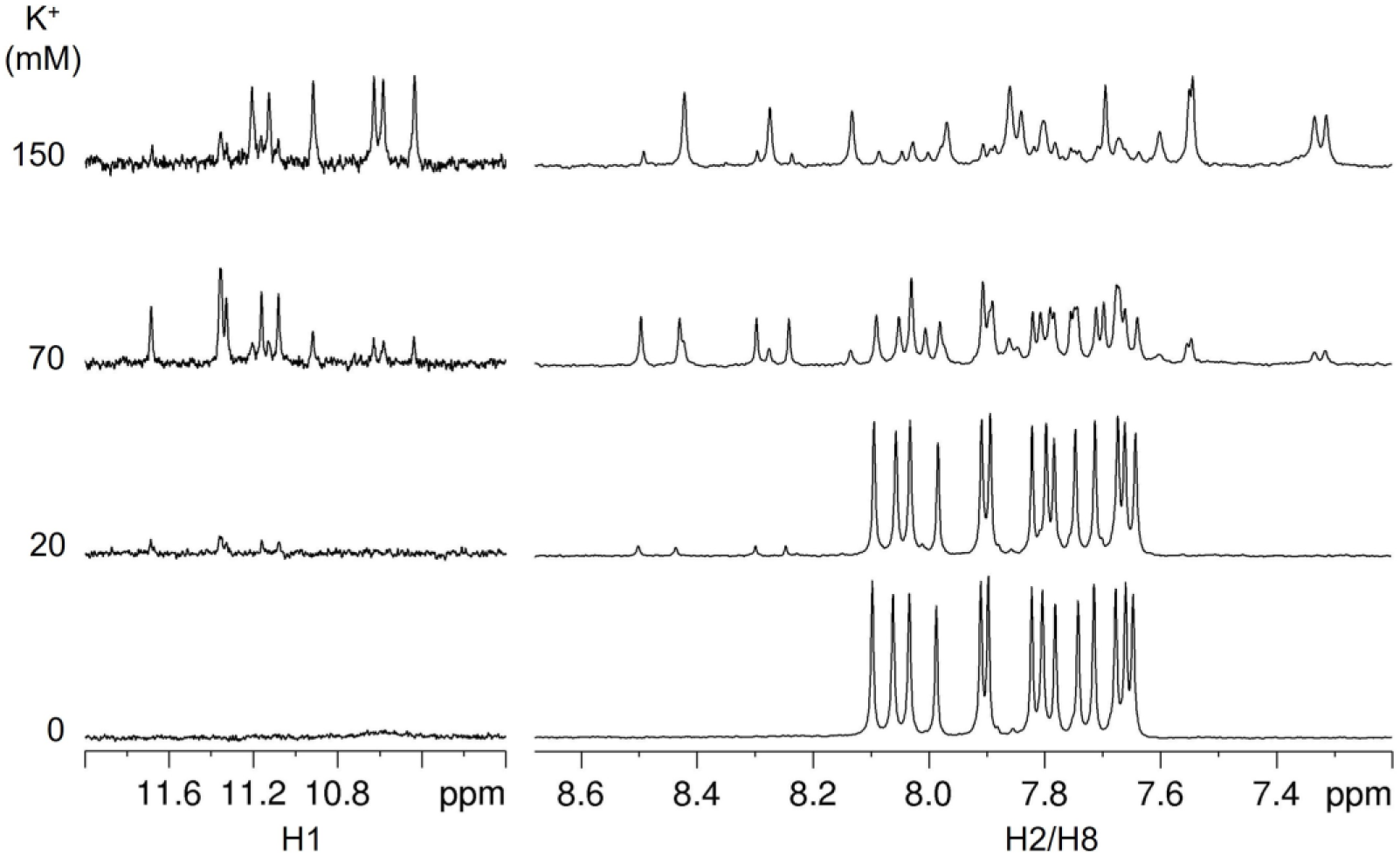
The 1D ^1^H NMR spectra (imino and aromatic proton regions) of d(AAGGG)_2_. [DNA] = 100 µM, [NaPi, pH 7] = 1 mM, [KCl] = 0/20/50/70/150 mM, 10% D_2_O, 25 °C.

**Fig. S2.**
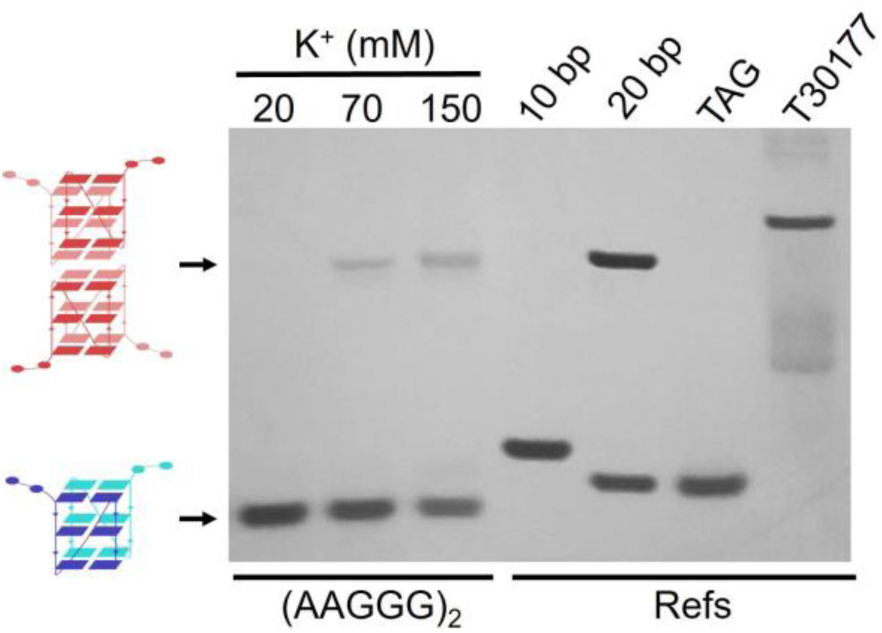
Native PAGE of d(AAGGG)_2_. [DNA] = 100 µM, [NaPi, pH 7] = 1 mM, 25 ℃. [KCl] = 20/70/150 mM for the d(AAGGG)_2_, [MgCl_2_] = 10 mM for the 10-bp and 20-bp duplex references, [KCl] = 90 mM for the reference of TAG that formed a three-layer dimeric G4 (12 nt × 2), and [KCl] = 70 mM for the reference of T30177 that formed a six-layer dimeric G4 (17 nt × 2).

**Fig. S3.**
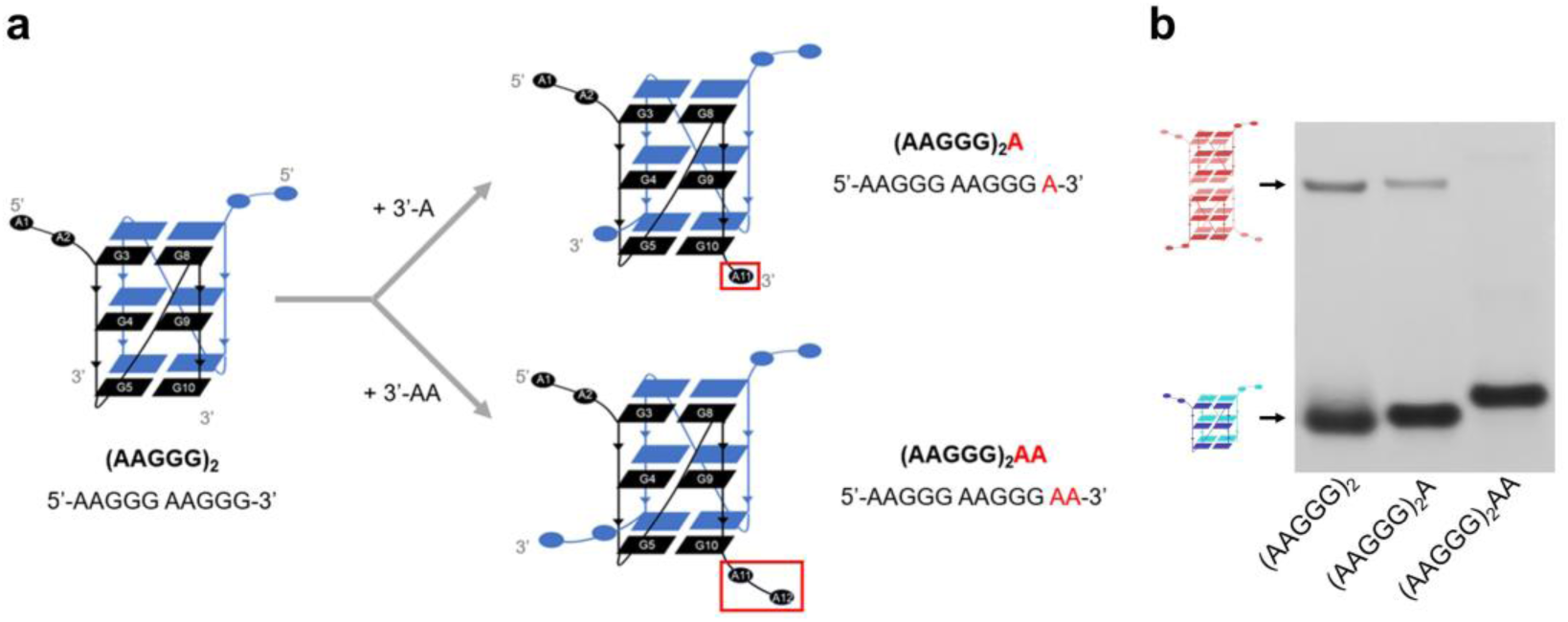
(a) Schematic of adding 3′-adenine residue(s) to d(AAGGG)_2_ to prevent the formation of tetrameric G4. (b) Native PAGE of d(AAGGG)_2_, d(AAGGG)_2_A and d(AAGGG)_2_AA. [DNA] = 100 µM, [NaPi, pH 7] = 1 mM, [KCl] = 150 mM, 25 ℃.

**Fig. S4.**
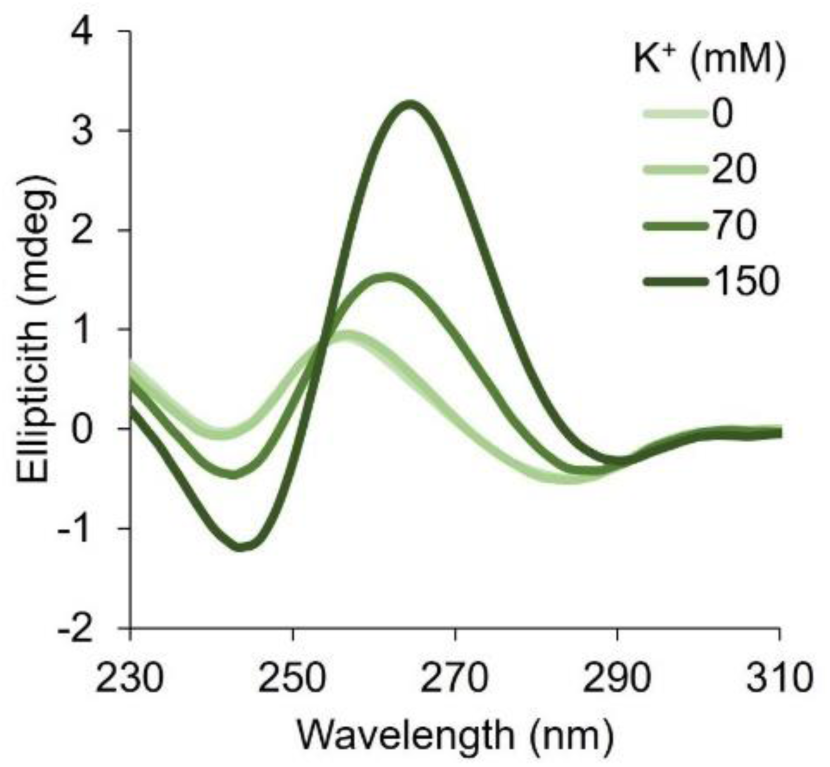
CD spectra of d(AAGGG)_2_AA. [DNA] = 20 µM, [NaPi, pH 7] = 1 mM, [KCl] = 0/20/70/150 mM, 25 °C.

**Fig. S5.**
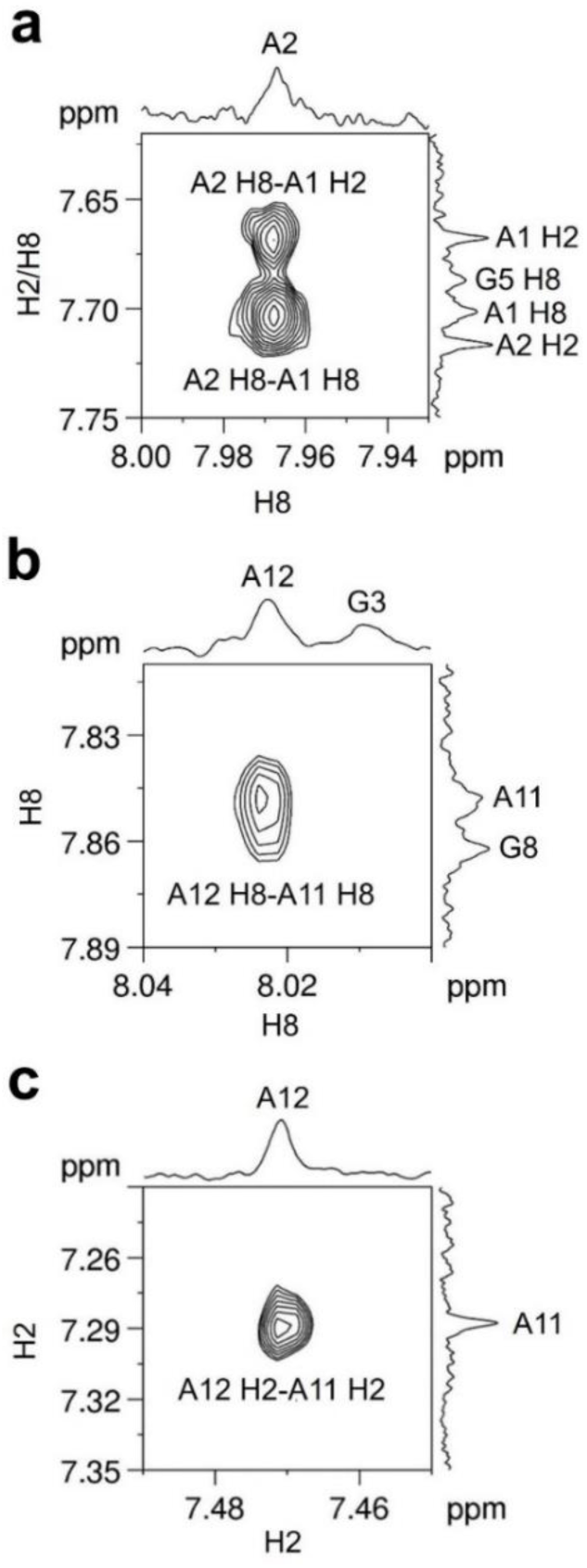
NOESY spectrum of d(AAGGG)_2_AA shows NOEs of (a) A2 H8-A1 H2 and A2 H8-A1 H8**, (b)** A12 H8-A11 H8 and (c) A12 H2-A11 H2. [DNA] = 800 µM, [NaPi, pH 7] = 1 mM, [KCl] = 150 mM, 99.96% D_2_O, 25 ℃.

**Fig. S6.**
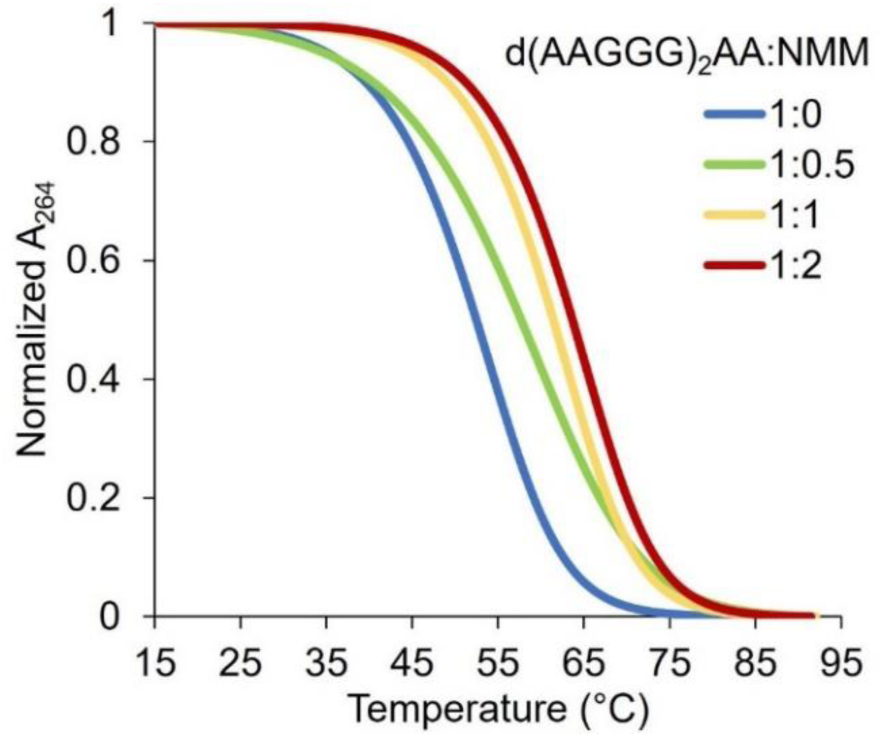
CD melting curves of d(AAGGG)_2_AA in various NMM concentrations. [DNA] = 20 µM, [NaPi, pH 7] = 1 mM, [KCl] = 150 mM.

**Fig. S7.**
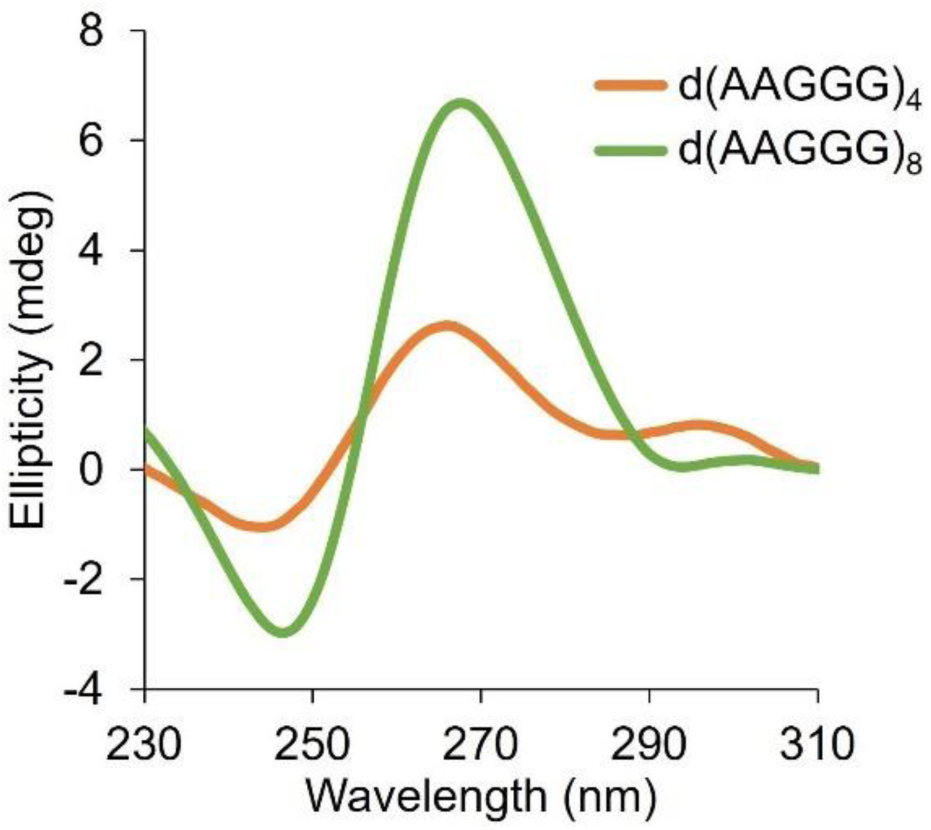
CD spectra of d(AAGGG)_4_ and d(AAGGG)_8_. [DNA] = 20 µM, [NaPi, pH 7] = 1 mM, [KCl] = 150 mM.

**Fig. S8.**
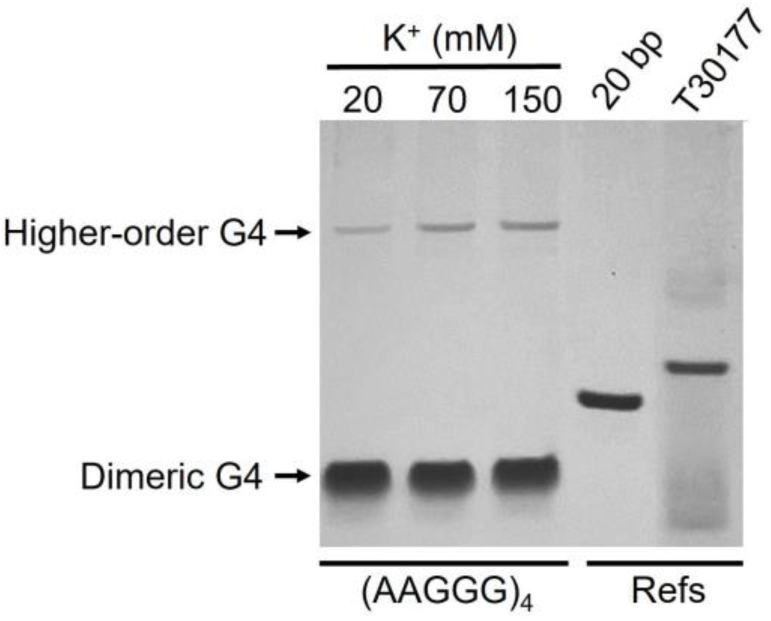
Native PAGE of d(AAGGG)_4_. [DNA] = 100 µM, [NaPi, pH 7] = 1 mM, 25 ℃. [KCl] = 20/70/150 mM for the d(AAGGG)_4_, [MgCl_2_] = 10 mM for the 20-bp duplex reference, and [KCl] = 70 mM for the reference of T30177 that formed a six-layer dimeric G4 (17 nt × 2).

## Notes

### Competing Interest Statement

The authors have declared no competing interest.

## References

1. Hannan, A.J. Tandem repeats mediating genetic plasticity in health and disease. Nat. Rev. Genet. 19, 286–298 (2018).

2. La Spada, A.R. & Taylor, J.P. Repeat expansion disease: progress and puzzles in disease pathogenesis. Nat. Rev. Genet. 11, 247–258 (2010).

3. Mirkin, S.M. Expandable DNA repeats and human disease. Nature 447, 932–940 (2007).

4. Cortese, A. et al. Cerebellar ataxia, neuropathy, vestibular areflexia syndrome due to RFC1 repeat expansion. Brain 143, 480–490 (2020).

5. Tranchant, C. & Anheim, M. CANVAS: a very late onset cerebellar ataxia, due to biallelic expansions in the RFC1 gene. Rev. Neurol. (Paris) 175, 493–494 (2019).

6. Cortese, A. et al. Biallelic expansion of an intronic repeat in RFC1 is a common cause of late-onset ataxia. Nat. Genet. 51, 649–658 (2019).

7. Dominik, N., Galassi Deforie, V., Cortese, A. & Houlden, H. CANVAS: a late onset ataxia due to biallelic intronic AAGGG expansions. J. Neurol. 268, 1119–1126 (2021).

8. Tsurimoto, T. & Stillman, B. Functions of replication factor C and proliferating-cell nuclear antigen: functional similarity of DNA polymerase accessory proteins from human cells and bacteriophage T4. PNAS 87, 1023–1027 (1990).

9. Rafehi, H. et al. Bioinformatics-based identification of expanded repeats: A non-reference intronic pentamer expansion in RFC1 causes CANVAS. Am. J. Hum. Genet. 105, 151–165 (2019).

10. Ronco, R. et al. Truncating variants in RFC1 in Cerebellar Ataxia, Neuropathy, and Vestibular Areflexia Syndrome. Neurology 100, e543–e554 (2023).

11. Benkirane, M. et al. RFC1 nonsense and frameshift variants cause CANVAS: clues for an unsolved pathophysiology. Brain 145, 3770–3775 (2022).

12. da Silva Schmitt, G., et al. Dopa-responsive parkinsonism in a patient With homozygous RFC1 Expansions. Mov. Disord. 35, 1889–1890 (2020).

13. Kytovuori, L. et al. Biallelic expansion in RFC1 as a rare cause of Parkinson’s disease. NPJ Parkinsons Dis. 8, 6 (2022).

14. Montaut, S. et al. Biallelic RFC1-expansion in a French multicentric sporadic ataxia cohort. J. Neurol. 268, 3337–3343 (2021).

15. Sullivan, R. et al. RFC1 intronic repeat expansions absent in pathologically confirmed Multiple Systems Atrophy. Mov. Disord. 35, 1277–1279 (2020).

16. Wan, L. et al. Biallelic intronic AAGGG expansion of RFC1 is related to Multiple System Atrophy. Ann. Neurol. 88, 1132–1143 (2020).

17. Tagliapietra, M. et al. RFC1 AAGGG repeat expansion masquerading as Chronic Idiopathic Axonal Polyneuropathy. J. Neurol. 268, 4280–4290 (2021).

18. Curro, R. et al. RFC1 expansions are a common cause of idiopathic sensory neuropathy. Brain 144, 1542–1550 (2021).

19. Bochman, M.L., Paeschke, K. & Zakian, V.A. DNA secondary structures: stability and function of G-quadruplex structures. Nat. Rev. Genet. 13, 770–780 (2012).

20. Varshney, D., Spiegel, J., Zyner, K., Tannahill, D. & Balasubramanian, S. The regulation and functions of DNA and RNA G-quadruplexes. Nat. Rev. Mol. Cell Biol. 21, 459–474 (2020).

21. Tu, J. et al. Direct genome-wide identification of G-quadruplex structures by whole-genome resequencing. Nat. Commun. 12, 6014 (2021).

22. Wang, E., Thombre, R., Shah, Y., Latanich, R. & Wang, J. G-quadruplexes as pathogenic drivers in neurodegenerative disorders. Nucleic Acids Res. 49, 4816–4830 (2021).

23. Tateishi-Karimata, H. & Sugimoto, N. Roles of non-canonical structures of nucleic acids in cancer and neurodegenerative diseases. Nucleic Acids Res. 49, 7839–7855 (2021).

24. Haeusler, A.R. et al. C9orf72 nucleotide repeat structures initiate molecular cascades of disease. Nature 507, 195–200 (2014).

25. Tseng, Y.J. et al. The RNA helicase DHX36-G4R1 modulates C9orf72 GGGGCC hexanucleotide repeat-associated translation. J. Biol. Chem. 297, 100914 (2021).

26. Liu, H. et al. A helicase unwinds hexanucleotide repeat RNA G-Quadruplexes and facilitates repeat-associated non-AUG translation. J. Am. Chem. Soc. 143, 7368–7379 (2021).

27. Phan, A.T. & Patel, D.J. Two-repeat human telomeric d(TAGGGTTAGGGT) sequence forms interconverting parallel and antiparallel G-quadruplexes in solution: distinct topologies, thermodynamic properties, and folding/unfolding kinetics. J. Am. Chem. Soc. 125, 15021–15027 (2003).

28. Mukundan, V.T., Do, N.Q. & Phan, A.T. HIV-1 integrase inhibitor T30177 forms a stacked dimeric G-quadruplex structure containing bulges. Nucleic Acids Res. 39, 8984–8991 (2011).

29. Wang, Z.F., Li, M.H., Hsu, S.T. & Chang, T.C. Structural basis of sodium-potassium exchange of a human telomeric DNA quadruplex without topological conversion. Nucleic Acids Res. 42, 4723–4733 (2014).

30. Sannapureddi, R.K.R., Mohanty, M.K., Salmon, L. & Sathyamoorthy, B. Conformational plasticity of parallel G-quadruplex horizontal line implications on duplex-quadruplex motifs. J. Am. Chem. Soc. 145, 15370–15380 (2023).

31. Tian, T., Chen, Y.-Q., Wang, S.-R. & Zhou, X. G-quadruplex: a regulator of gene expression and its chemical targeting. Chem. 4, 1314–1344 (2018).

32. Kreig, A. et al. G-quadruplex formation in double strand DNA probed by NMM and CV fluorescence. Nucleic Acids Res. 43, 7961–7970 (2015).

33. Madabhushi, R., Pan, L. & Tsai, L.-H. DNA damage and its links to neurodegeneration. Neuron. 83, 266–282 (2014).

34. Jeppesen, D.K., Bohr, V.A. & Stevnsner, T. DNA repair deficiency in neurodegeneration. Prog. Neurobiol. 94, 166–200 (2011).

35. Malik, I., Kelley, C.P., Wang, E.T. & Todd, P.K. Molecular mechanisms underlying nucleotide repeat expansion disorders. Nat. Rev. Mol. Cell Biol. 22, 589–607 (2021).

36. Robinson, J., Raguseo, F., Nuccio, S.P., Liano, D. & Di Antonio, M. DNA G-quadruplex structures: more than simple roadblocks to transcription? Nucleic Acids Res. 49, 8419–8431 (2021).

37. Takahashi, S., Brazier, J.A. & Sugimoto, N. Topological impact of noncanonical DNA structures on Klenow fragment of DNA polymerase. Proc. Natl. Acad. Sci. U S A 114, 9605–9610 (2017).

38. Guo, P. & Lam, S.L. Minidumbbell structures formed by ATTCT pentanucleotide repeats in spinocerebellar ataxia type 10. Nucleic Acids Res. 48, 7557–7568 (2020).

39. Viguera, E., Canceill, D. & Ehrlich, S.D. Replication slippage involves DNA polymerase pausing and dissociation. EMBO J. 20, 2587–2595 (2001).

40. Khristich, A.N. & Mirkin, S.M. On the wrong DNA track: molecular mechanisms of repeat-mediated genome instability. J. Biol. Chem. 295, 4134–4170 (2020).

41. Lyu, K., et al. An RNA G-quadruplex structure within the ADAR 5’UTR interacts with DHX36 helicase to regulate translation. Angew. Chem. Int. Ed. Engl. 61, e202203553 (2022).

42. Amrane, S. et al. Deciphering RNA G-quadruplex function during the early steps of HIV-1 infection. Nucleic Acids Res. 50, 12328–12343 (2022).

43. Dumetz, F. et al. G-quadruplex RNA motifs influence gene expression in the malaria parasite Plasmodium falciparum. Nucleic Acids Res. 49, 12486–12501 (2021).

44. Jain, A. & Vale, R.D. RNA phase transitions in repeat expansion disorders. Nature 546, 243–247 (2017).

45. Fay, M.M., Anderson, P.J. & Ivanov, P. ALS/FTD-associated C9ORF72 repeat RNA promotes phase transitions in vitro and in cells. Cell Rep. 21, 3573–3584 (2017).

46. Tang, F., Wang, Y., Gao, Z., Guo, S. & Wang, Y. Polymerase eta recruits DHX9 helicase to promote replication across guanine quadruplex structures. J. Am. Chem. Soc. 144, 14016–14020 (2022).

47. Liano, D., Chowdhury, S. & Di Antonio, M. Cockayne syndrome B protein selectively resolves and interact with intermolecular DNA G-Quadruplex structures. J. Am. Chem. Soc. 143, 20988–21002 (2021).

48. Hosur, R.V., Govil, G. & Miles, H.T. Application of two-dimensional NMR spectroscopy in the determination of solution conformation of nucleic acids. Magn. Reson. Chem. 26, 927–944 (1988).

49. D.A. Case, et al. AMBER 2018. (University of California, San Francisco; 2018).

50. Ivani, I. et al. Parmbsc1: a refined force field for DNA simulations. Nat. Methods 13, 55–58 (2016).

51. Greenfield, N.J. Using circular dichroism collected as a function of temperature to determine the thermodynamics of protein unfolding and binding interactions. Nat. Protoc. 1, 2527–2535 (2006).

